# Layer-by-Layer Fabrication of 3D Hydrogel Structures Using Open Microfluidics

**DOI:** 10.1101/687251

**Authors:** Ulri N. Lee, John H. Day, Amanda J. Haack, Wenbo Lu, Ashleigh B. Theberge, Erwin Berthier

**Affiliations:** University of Washington, Department of Chemistry, Seattle, WA; University of Washington School of Medicine, Medical Scientist Training Program, Seattle, WA; University of Washington School of Medicine, Department of Urology, Seattle, WA

**Keywords:** Layer-by-layer patterning, 3D fabrication, hydrogels, open microfluidics, capillary flow

## Abstract

Patterning and 3D fabrication techniques have enabled the use of hydrogels for a number of applications including microfluidics, sensors, separations, and tissue engineering in which form fits function. Devices such as reconfigurable microvalves or implantable tissues have been created using lithography or casting techniques. Here, we present a novel open microfluidic patterning method that utilizes surface tension forces to pattern hydrogel layers on top of each other, producing 3D hydrogel structures. We use a patterning device to form a temporary open microfluidic channel on an existing gel layer, allowing the controlled flow of unpolymerized gel in regions defined by the device. Once the gel is polymerized, the patterning device can then be removed, and subsequent layers added to create a multi-layered 3D structure. The use of open-microfluidic and surface tension-based methods to define the shape of each layer enables patterning to be performed with a simple pipette, minimizing dead-volume and shear stress applied on the fluid. Our method is compatible with unmodified (native) biological hydrogels, or other non-biological materials with fluid properties compatible with capillary flow. With our open-microfluidic layer-by-layer fabrication method, we demonstrate the capability to build agarose and type I collagen structures featuring asymmetric designs, multiple components, overhanging features, and cell laden regions.

## INTRODUCTION

Hydrogels are configurable and functional materials that can be patterned to obtain desired functions ranging from mechanical responses (e.g., microfluidic valves and pumps) to organotypic functions (e.g., biological models and implantable materials). Many of these applications have simple patterning requirements, such integration in microfluidic devices as valves,^1, 2^ for encapsulating colorimetric indicators,^3^ or for chemical separation of desired analytes.^4, 5^ For cell culture applications, hydrogel patterning techniques are used for encapsulating cells in a controlled position relative to additional cell types, microchannels or extracellular matrix components.^6-14^ Reliable and low-entrance barrier solutions to these applications would have the potential of widespread adoption in biology, fluidics, and engineering labs that seek prototyping solutions. Here, we present a method for multilayer hydrogel patterning in three dimensions that uses a simple setup: patterning devices are positioned above a substrate to guide the flow of hydrogels using surface tension forces. Hydrogels are delivered with standard pipettes, minimizing dead volume, a key consideration when patterning expensive hydrogels or hydrogels laden with rare cell types.

Many researchers cannot easily use current hydrogel patterning methods due to the investment in time or skills required, equipment cost, or the constraints on materials or cell types. Amongst these methods, photopolymerization techniques are robust and offer resolution on the order of microns,^1, 6, 10, 12, 13, 15-19^ however photo-initiators have been shown to cause cellular damage,^20, 21^ and the necessity for chemical functionalization of the gel precludes the use of native (unmodified) gels. Other broadly used hydrogel patterning methods include inkjet^22-24^ and microextrusion^25-29^ printing. In these methods, the hydrogel is jetted through a nozzle that exposes any cells encapsulated within the hydrogel to shear stress, which can mechanically damage the cells and limit the types of cells that can be patterned.^30-32^ Inkjet and extrusion based printing methods can also require optimization depending on the viscosity and type of gel used. Many of these hydrogel patterning techniques can require large volumes of fluid to fill vats, fluidic channels, or tubing used in typical gel 3D printing setups, limiting the use of rare cell types or expensive gels. Casting hydrogels using a fabricated negative mold provides an alternative approach, but the complexity of the gel structures obtainable is limited due to demolding considerations.^7, 33-35^ Therefore, a hydrogel patterning method that does not rely on photoinitiators, external pressure, tubing, or reservoirs of gel could reduce material waste and alleviate some of the stresses experienced by cells, opening up the potential to use more fragile, primary, or rare cell types.

In the present manuscript, we present a novel layer-by-layer patterning method that uses open microfluidic principles and surface tension driven flow to pattern sequential hydrogel layers, each delivered directly using a pipette. To pattern each hydrogel layer, we use a patterning device, which is either a plastic physical ‘rail’ or a hydrophilic track delineated on a hydrophobic flat surface; Both enable the pre-gel solution to flow via capillary action without external pressure and constrain the pre-gel solution under the rail footprint via capillary pinning.^36, 37^ In this work, ‘patterning device’ refers to either a rail or a hydrophobic flat surface with a hydrophilic track. The patterning device is placed above an existing hydrogel layer at a designed height. The structures in this work were made with layer heights between 100-500 *µ*m, although the method is not limited to heights within this range. The gap between the patterning device and the existing hydrogel layer forms an open microfluidic ‘channel’ in which liquid pre-gel solution can flow, where the channel ‘ceiling’ is the patterning device, the ‘floor’ is the previous gel layer, and the ‘sides’ are free air-liquid interfaces (Figure 1). The patterning device can be removed once the pre-gel solution is gelled, and another patterning device can be applied to pattern the next layer, providing the ability to build three-dimensional structures layer-by-layer. This work builds on prior publications demonstrating the use of rails to pattern a single hydrogel layer to partition a well plate for coculture^37-40^ where due to the design constraints of the previous rail devices, the patterning device could not be removed once the hydrogel was polymerized, and thus a three dimensional hydrogel structure could not be fabricated. Here, we built multilayer hydrogel structures from native hydrogels which included unsupported overhanging features, objects comprising multiple materials and components both within a layer and across layers, asymmetric designs, and a cell-laden gel architecture. The ability to establish a full layer in a single dispensing step is distinct from extrusion or inkjet patterning methods, which require multiple passes of material deposition for a single layer. Further, in contrast to methods that use photopolymerizable gels and light as a means of patterning hydrogels, which then require washing away of unused, pre-gel solution,^1, 6, 16, 17^ we limit waste, as all of the pre-gel solution that is dispensed is polymerized and part of the desired pattern.

**Figure 1.**
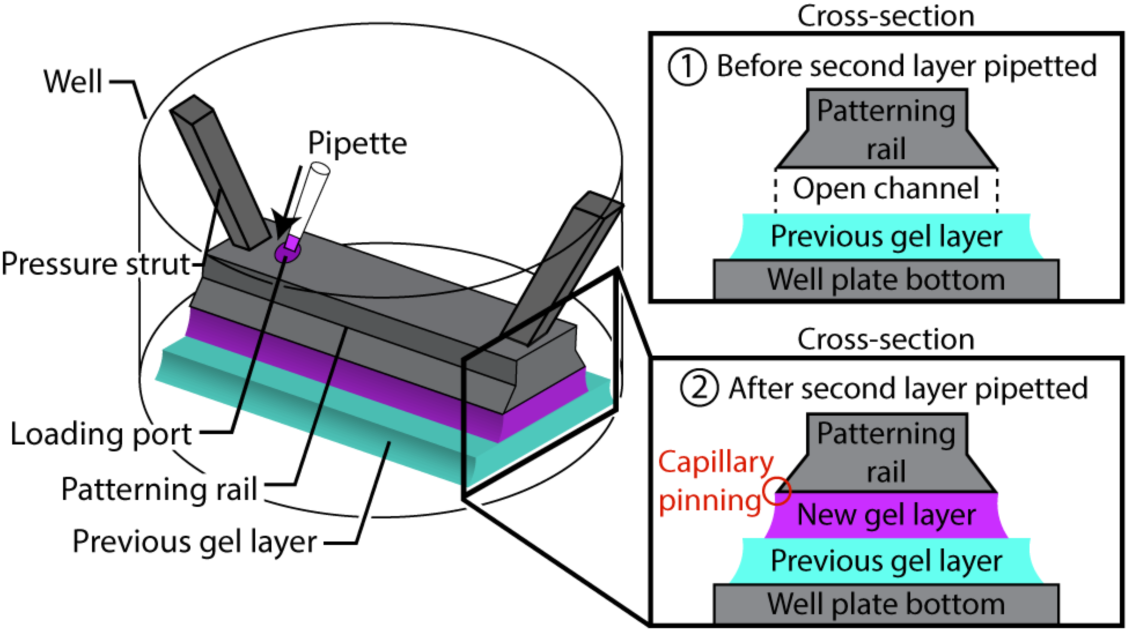
Schematic showing the mechanism of layer-by-layer gel patterning using open microfluidics. An open channel is created above an existing gel layer using a rail-based patterning device.

## EXPERIMENTAL

### Patterning Device Design and Fabrication (Figures 2-6)

Patterning devices were designed in Solidworks 2017 and 3D printed out of clear resin (RS-F2-GPCL-04) using a Form 2 stereolithography 3D printer (Formlabs) (Figures 2-6) or milled (Figure 6) on a computer numerical control (CNC) mill (PCNC770, Tormach). Design files for the patterning devices in Figure 2B and the template used as the base for all patterning devices are included in the Supporting Information. The patterning devices were sonicated in isopropyl alcohol (IPA) for 10 minutes and again in clean IPA for 5 minutes to remove excess uncured resin. The devices were dried with compressed air and cured under a 395-405 nm 20W UV lamp (Quans) for 1 hour.

**Figure 2.**
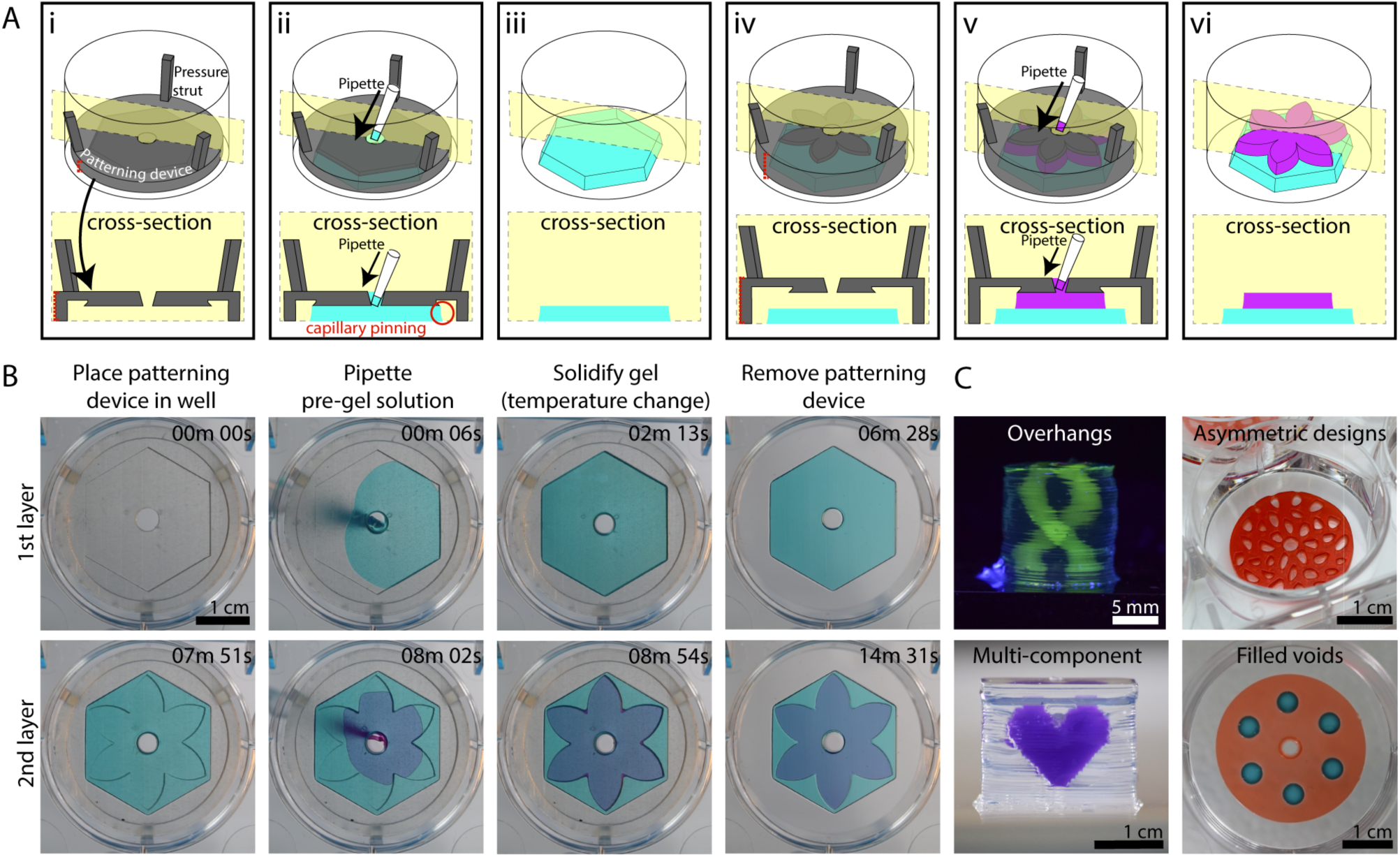
Rail-based open microfluidic layered patterning of unmodified hydrogels. (A) Schematic of patterning two layers of gel. (Ai) Patterning device (rail) is placed in a 6-well plate; the patterning region sits 250 *µ*m above the well plate surface. Red dotted line on the left indicates portion of patterning device that is adjusted to determine the thickness of the gel layer. (Aii) Pre-gel solution is pipetted into the pipette port. (Aiii) Patterning device is removed after the pre-gel solution is solidified. (Aiv-Avi) Second layer is patterned using the same steps for the first layer. Red dotted line on the left in Aiv indicates distance between the well plate and the rail after an initial layer of gel is patterned. Note that the dotted line is longer than the red line seen in Ai. Cross-sections of patterning devices show capillary pinning (red circle) and 45° angle around the edges of the device to prevent capillary rise along the vertical face of the patterning device. A magnified schematic is included in Figure S1; an engineering drawing of the hexagon and petal devices is included in Figure S2. (B) Images of the patterning process for 1.5% low gelling temperature agarose dyed for visualization with teal and magenta India ink. Video was recorded from the underside of a 6-well plate. Full video available in the SI. (C) Various agarose structures built using rail-based open microfluidics to showcase the power of the method. Cylindrical structure with hollow unsupported winding tubes (double helix) (top, left), asymmetric geometries patterned in a well plate (top, right), a multicomponent structure (bottom, left), and a single layer of gel with islands of voids filled with a second gel in a single pipetting step (bottom, right).

**Figure 3.**
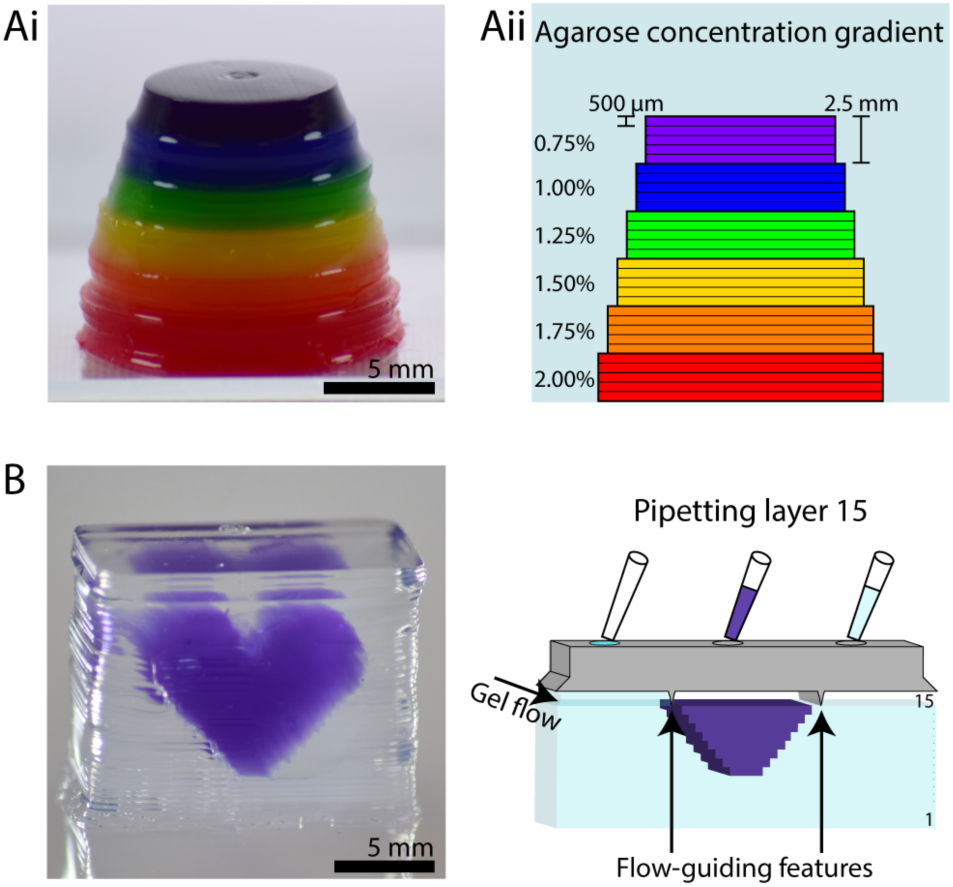
Constructing heterogeneous structures. (A) Structure with multiple agarose concentrations. (Ai) Photo of agarose structure (dyed for visualization, with each color representing a different concentration) in ascending layers. Layers are 500 *µ*m thick, and each color contains 5 layers. (Aii) Cross-sectional schematic showing the agarose concentrations in each layer. (B) Purple agarose heart inside a colorless agarose cube demonstrating the ability to pattern materials of different composition *within* a layer (left). Schematic of patterning layer 15 of the 34-layered structure with compartmentalization features to segregate the different gels (right). Layers are 500 *µ*m thick.

**Figure 4.**
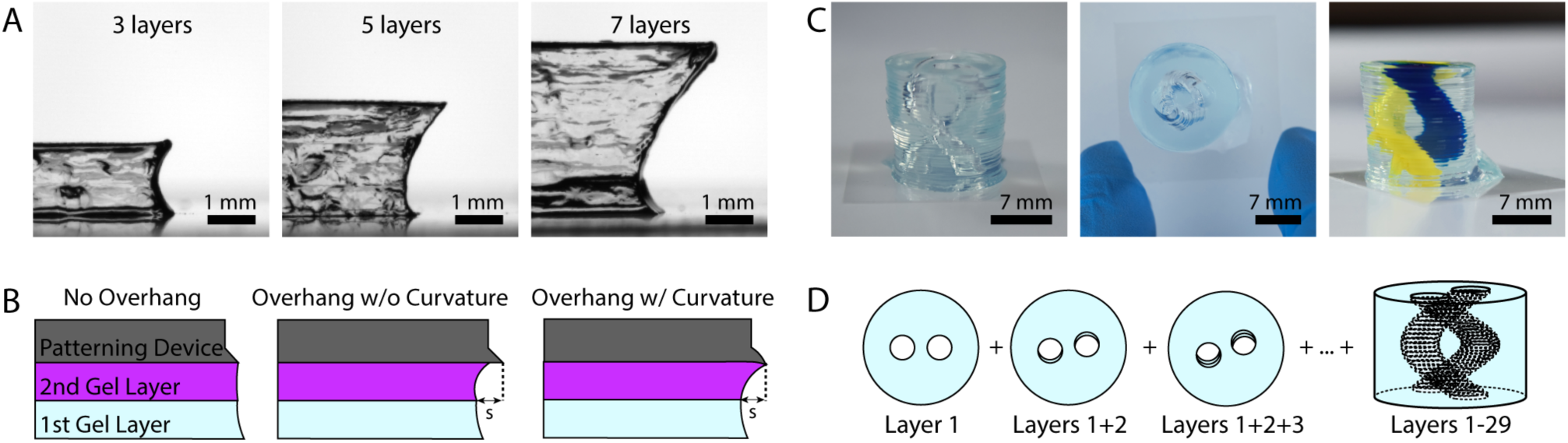
Rail-based open microfluidic layered patterning of overhanging features. (A) Imaging of a cross-section of an overhanging agarose structure at three different points of production, 3 layers (left), 5 layers (center) and 7 layers (right). (B) Schematic for rail design of overhanging features showing no overhang, an overhang without curvature and an overhang with downward curvature; the latter rail design was used to print the structures shown in A and C. (C) Images of cylindrical structure with hollow unsupported winding tubes (double helix) made from agarose, showing side view (left), top down image (center), and side view where tubes are filled with yellow and blue India ink dyes for visualization (right). Full description of design in the Supporting Information and Figure S3. (D) Schematic showing the process for building the double helix structure, where the blue color represents the patterned agarose. Each layer is rotated about the center from the previous layer, forming a hollow helix within a cylinder of agarose.

**Figure 5.**
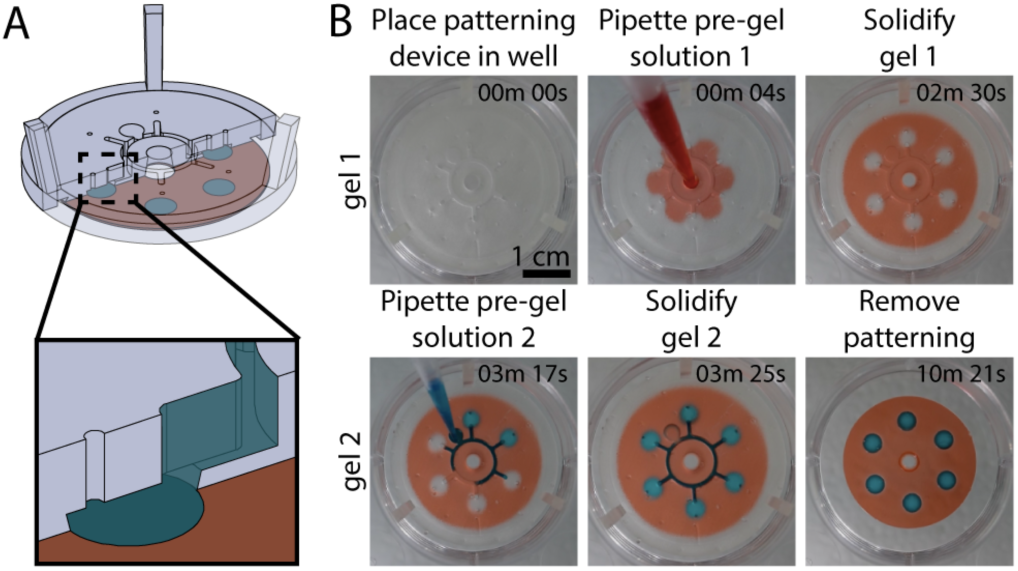
Integrated fluidic channels in patterning device for depositing second material in a layer. (A) Cross-sectional schematic representation of ‘void-filling’ patterning device with underlying layer. (B) Video stills of patterning red colored agarose (250 *µ*m thick) and filling voids with blue colored agarose (750 *µ*m thick). Full video available in the SI.

**Figure 6.**
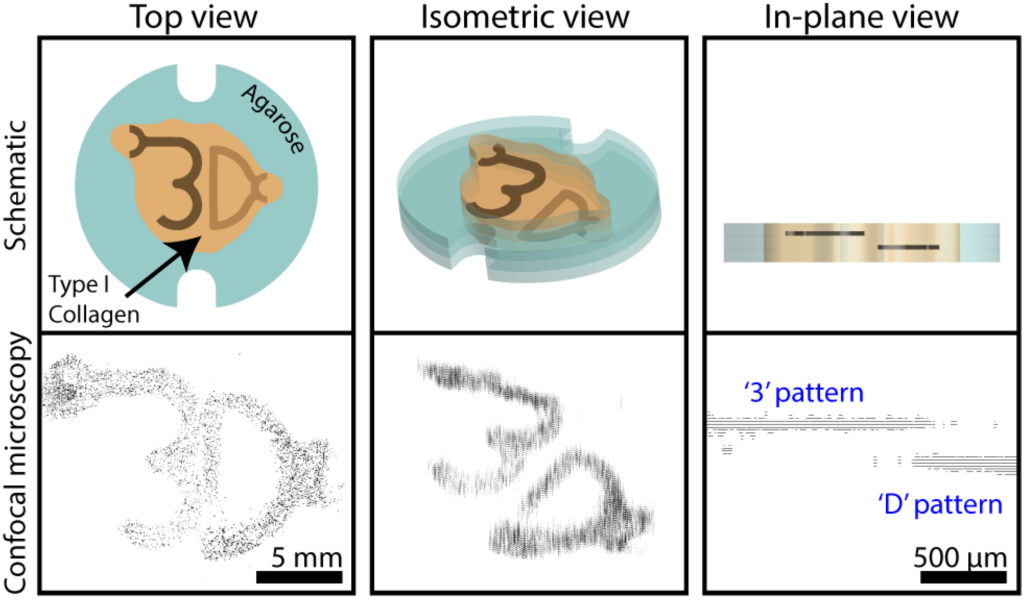
Schematics and confocal images of human fetal lung fibroblasts in a type I collagen structure. Cells were stained with Calcein AM (shown in black). The top view comprises 130 fields of view (which each contain 27 Z-stacked images) stitched together. Cells that make up the “3” and “D” are each printed in separate 100 *µ*m thick collagen layers that are separated from each other by a 200 *µ*m thick cell-free collagen layer. Z-step value was increased tenfold in isometric view for visualization purposes. Original confocal image file is included in the Supporting Information in Figure S5. Schematic of the workflow used to achieve this structure is shown in Figure S4; note: the additional features on the top left of the “3” and the right of the “D” were an intentional part of the design, used as loading ports for the pre-gel solution.

### Plasma Treating Patterning Devices (Figures 2-7)

**Figure 7.**
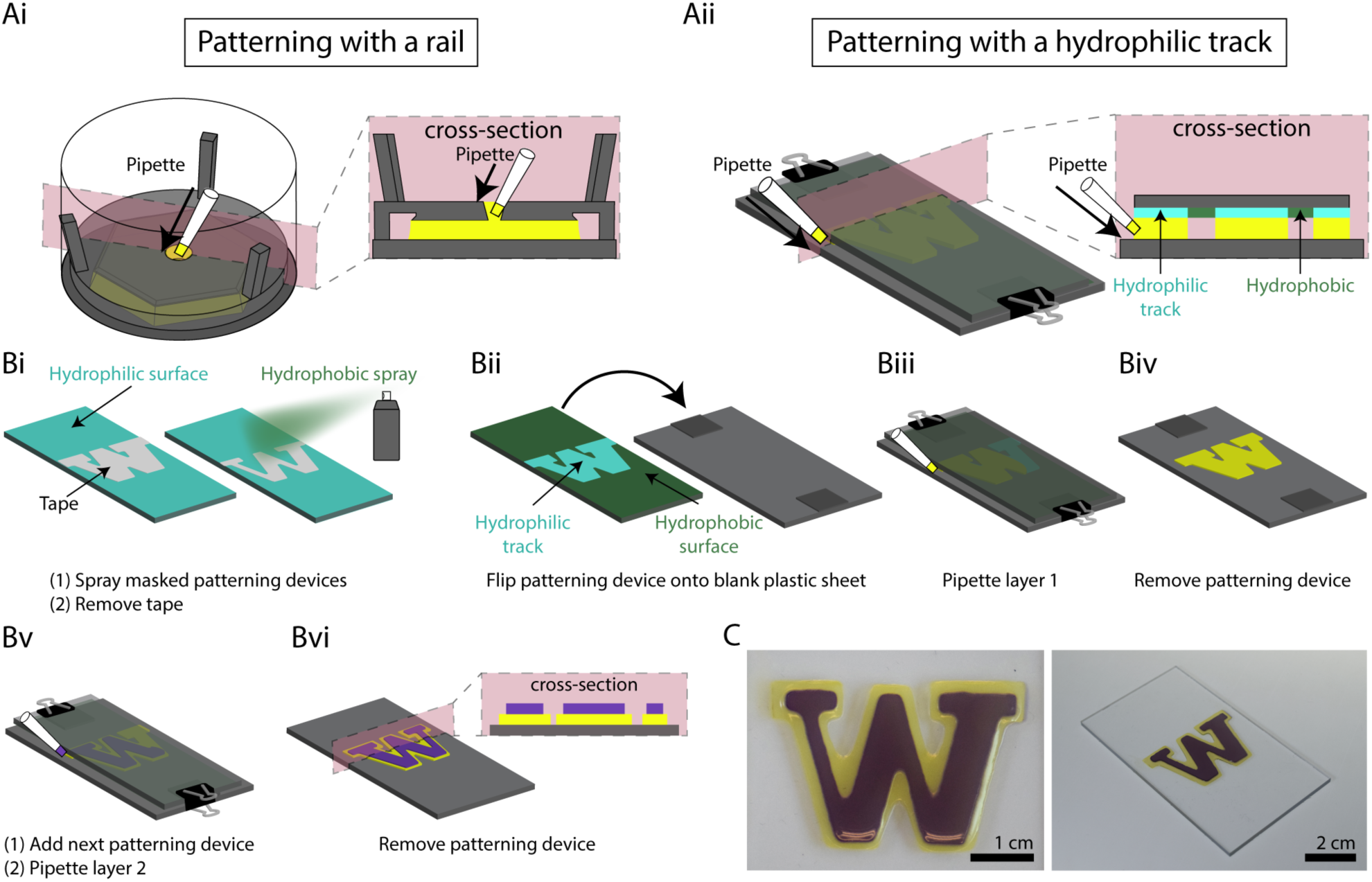
Layer-by-layer hydrogel patterning without a rail. (A) Overview of two patterning processes contrasting (Ai) patterning with a rail and (Aii) patterning with hydrophilic tracks. (Bi) Plasma treated polystyrene (PS) sheets were masked with tape, and a hydrophobic spray was applied. After the hydrophobic layer had set, the tape was removed. (Bii) The hydrophobic side of the PS sheet placed face down onto a clean, plasma treated PS sheet with stacked coverslips as spacers to create a gap between the two sheets. (Biii) The patterning device was secured to the spacers and bottom substrate with binder clips and the pre-gel solution was pipetted in the gap between the patterning device and bottom substrate at both sides of the ‘W’. (Biv) The patterning device was removed after the gel solidified. (Bv) Additional coverslips were stacked on the existing spacers to create a larger gap for the second layer, and the second patterning device was aligned by eye, placed and secured with binder clips. The second layer of pre-gel solution was then pipetted in the same way. (Bvi) The patterning device was removed to reveal the 2-layered hydrogel structure. The thickness of each layer was 500 *µ*m (C) Top view (left) and isometric (right) photo of two-layered agarose ‘W’. Video of patterning process is available in the Supporting Information.

The patterning devices were plasma treated for increased wettability using a Zepto plasma treater (Diener Electronic). The chamber was pumped down to a pressure of 0.20 mbar, gas (free air) was supplied (4 min, 0.25 mbar), and power enabled (2 min, 200 W).

#### Agarose Gel Composition

Low gelling temperature agarose (MilliporeSigma) was dissolved in deionized water to a concentration of 15 mg/mL and dyed for visual aid with food dye (Spice Supreme), fluorescein labeled dextran (70,000 MW, Invitrogen), or India ink (Dr. Ph Martin’s). All agarose structures used 1.5% wt/v gel except for the multicomponent structure in Figure 3A.

### Fabrication of Overhanging Structures (Figure 4)

For the seven layer overhang structure, each layer height was 500 *µ*m. Layers 2 and 3 were designed to be 239.9 *µ*m of overhang, whereas the following layers were designed to be 479 *µ*m of overhang (*s* in Figure 4B). A schematic of the patterning device design and detailed dimensions can be found in the Supporting Information.

### Fabrication of Hollow Double Helix (Figure 4)

Using the patterning device in a 6-well plate, each layer was allowed to set at room temperature for approximately 10-20 minutes prior to removal of the patterning rail. Each helix was constructed over 2-3 days; the total fabrication time if done at once was about 10 hours. Sacrificial water was added to the empty wells in the 6-well plates to prevent drying when stored overnight. More details on the design of each layer can be found in the Supporting Information.

### Cell Culture of Human Fetal Lung Fibroblasts (Figure 6)

Agarose prepared for cell culture experiments was dissolved in 1X PBS at 1.5% wt/v and autoclaved. Type I collagen solutions were neutralized by mixing rat tail collagen (Corning #354249) with 500 mM HEPES in a 9:1 ratio, respectively. Cell suspensions containing human fetal lung (HFL) fibroblasts (ATCC) at concentrations of ∼9 million cells/mL were then added to the collagen solutions in a 1:2 ratio, respectively, to give a final collagen concentration of 5 mg/mL and a final cell concentration of ∼3 million cells/mL. Cell-laden structures were incubated at 37 °C for 24 hours in Dulbeco’s modified eagle medium (Gibco) prior to imaging. Cells were stained with Calcein AM and ethidium homodimer (MilliporeSigma). Cell-laden structures were imaged with a Leica TCS SP5 II confocal microscope and a 17.9 *µ*m Z-step. Images were processed in FIJI software.

### Patterning Type I Collagen (Figure 6)

Patterning collagen was achieved with the addition of a flattening step (Figure S4). This step involves placing a patterning device over the collagen layer prior to gelling such that the collagen will wet the pattern on the device, creating a flat surface to guide flow for the next layer. When the collagen is gelled and the device removed, the height of the resulting gel layer is precisely set in the shape and location of the pattern to be flowed in the next layer.

### Hydrophobic coating for patterning (Figure 7)

Rectangular polystyrene (PS) pieces were cut using a CNC mill (DATRON neo) to the same width as the designed “W” pattern. All PS pieces were plasma treated using the same protocol for the 3D printed patterning devices. The ‘W’ pattern, was cut from masking tape (3M) using a plotter cutter (Graphtec CE6000-40) and then applied to the plasma treated side of the PS, such that the edges of the ‘W’ lined up with the edges of the PS. Two coats of Step 1 of Rust-Oleum Neverwet™ hydrophobic treatment were applied, allowed to dry for 10 minutes, and then three coats of Step 2 were applied, and left to dry for an additional 10 minutes. The masking tape was removed, revealing a hydrophilic pattern with hydrophobic surroundings.

### Patterning hydrogels using hydrophilic/hydrophobic interfaces (Figure 7)

Two No. 1.5 coverslips (Fisherbrand) were stacked on top of a PS base at both ends. Double-sided tape was placed between each coverslip to prevent slipping during assembly. The total height of the coverslips and double-sided tape was measured to be 500 *µ*m with a caliber. The patterning piece (described above in the preceding section) was placed facedown over the coverslips and base. Binder clips were used to secure the PS layers and coverslips. A picture of the setup for the first layer can be found in Figure S6. 1.5% w/v agarose was dyed purple and yellow using India Ink (Dr. Ph Martin’s). The first layer was patterned by placing the pipette at the edge of the hydrophilic patterning region in between the patterning piece and the base. Both ends of the W were used as inlets. The agarose was allowed to solidify for 10 minutes before the patterning device was removed. To achieve the second layer, two additional coverslips and double-sided tape were added to increase the distance between the base and the second patterning device, which was aligned by eye. The gel loading and setting process for the second layer was the same as for the first layer. A video of the patterning can be found in the Supporting Information.

### Image Processing

Images were processed using Adobe Photoshop CC 2018 using a uniform brightness/contrast adjustment.

## RESULTS AND DISCUSSION

Our layer-by-layer patterning method uses open microfluidic principles to create temporary open channels on a plastic or existing hydrogel surface allowing pre-gel solution to flow in a spatially controlled area defined by a patterning rail or hydrophilic track (Figure 1). The gel can then be polymerized to create a patterned layer of hydrogel tens to hundreds of micrometers thick. The input of gel is powered by spontaneous capillary flow (SCF), which describes the surface tension-based flow of fluids in channel geometries that can have any number of open-air interfaces along the fluid path.^36, 40^ The geometric control of the gel layer is obtained by abrupt changes in geometry (in the case of a patterning rail) or hydrophilicity (in the case of hydrophilic tracks) which prevent fluids from flowing past these capillary pinning points. Together, the principles of SCF and capillary pinning enable the creation of a patterned gel layer with simple laboratory equipment (e.g., pipettes) in existing laboratory platforms (e.g., well plates) without requiring actuators, electronics, or pressure sources.

Figure 2 shows a specific embodiment of the patterning technique where a patterning rail is used to define the outline of a virtual open microfluidic channel. The patterning rail is placed at the location where the gel will be patterned and sits tens to hundreds of micrometers over the surface of the material below. The geometrical shape of the patterning area is designed to enable the spontaneous capillary flow of pre-gel solution throughout the open channel made between the patterning rail and the surface of the material below; capillary pinning enables the pre-gel solution to remain in the defined pattern. The patterning devices can be fabricated in hard plastic, such as resin that is 3D printed from a stereolithography 3D printer or polystyrene (PS) sheets that are CNC milled. The patterning device shown in Figure 2 fits inside of a 6-well plate, but the method is not limited to a 6-well plate form. The device sits flush to the floor of the well plate with spacers to control the height of the gel layer being formed (dotted red line in Figure 2Ai and iv, Figure S1, Figure S2) and pressure struts to maintain concentric alignment of the rail within the well.

In this manuscript, we patterned 100-500 *µ*m thick gel layers; numerical simulations of SCF suggest thinner layers (reduced *h* in Eq. 1) are achievable and, in fact, make it easier to satisfy the condition for SCF in our system which is defined by:

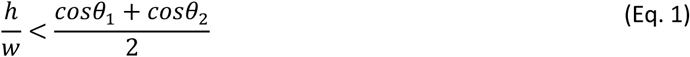

Where *h* is the height of the patterned layer (defined by the distance between the patterning area of the rail and the underlying substrate), *w* is the width of the rail perpendicular to the direction of flow, *θ*_1_ is the contact angle of the hydrogel on the patterning device, and *θ*_2_ is the contact angle of the hydrogel on the underlying substrate (conventionally equal to zero when flowing hydrogel over an existing hydrogel layer of the same type). To note, viscosity does not play a role in the condition for flow but only on the dynamics of the flow. A derivation of Eq. 1 from the generalized Cassie equation is provided in the Supporting Information.

When pre-gel solution is loaded into a patterning setup, it fills the ‘virtual channel’, and remains suspended between the patterning device and the underlying substrate until a polymerization trigger is introduced. Although temperature change was used for polymerization in this work, the ability to polymerize a layer after it has been patterned makes it possible to build multilayered structures of any hydrogel whose pre-gel solution can meet the condition for SCF. After a single layer of pre-gel solution is patterned and solidified (Figure 2Ai and 2Aii), the patterning device is removed (Figure 2Aiii), and a new patterning device is placed over the gel layer to define a new area for the next layer of gel (Figure 2Aiv). The patterning process is repeated for each additional layer (Figure 2Av and 2Avi). With this method we are able to deposit an entire layer in one pipetting step, as opposed to alternative nozzle-based 3D patterning methods which require multiple passes of material deposition for a single layer.

It is possible to pattern a pre-gel solution on top of an existing hydrogel layer using our method because capillary pinning still occurs at the junction of the edge of the patterning area and the previous hydrogel layer, maintaining a defined pattern (Figure 2B). However, gel overflow is observed for certain features; as seen in the video in the Supporting Information (corresponding to Figure 2B), overflow of dispensed fluid at the edges of the ‘flower’ patterning device occurs because the surface tension of the pre-gel solution, which tends to favor a rounded fluid front, overcomes pinning at the acute internal angles of the patterning device. Due to the manual nature of our method in its current form, applications requiring only a few layers are more suitable for immediate use of the technology. However, we also present several structures with multiple layers (>30) to demonstrate the ability of our layer-by-layer fabrication method to create three-dimensional structures. Figure 2C highlights some of these features we achieved with our layered patterning method which include unsupported overhangs, asymmetric geometries, multicomponent structures and cell-laden patterns. These structures are described in more detail in the subsequent figures.

The ability to pattern multiple materials in a single structure is of great interest for a variety of applications ranging from creating regions of varying material properties for studying molecular transport,^41, 42^ to controlling the gel porosity in specific regions of a microfluidic device for use in chemical separation assays.^4^ Patterning multiple materials is also of great importance to organotypic modelling. The cellular environment in biological tissue is heterogeneous in nature, and recapitulating this environment requires spatial control of cells in relation to multicomponent hydrogels and extracellular matrix materials. Our open microfluidic layered patterning method presents a new approach to building multicomponent structures. Figure 3A shows how different materials can be introduced at every layer to create a structure with differential material properties in the Z-direction. In Figure 3A, an agarose structure was erected beginning with a concentration of 2% wt/v and decreasing 0.25% wt/v every five layers to a final concentration of 0.75% wt/v. Each concentration is dyed a different color for visualization. The resulting structure has a stepwise concentration gradient in the Z-direction. The agarose concentrations were chosen here as a proof of concept to demonstrate the ability to pattern different compositions across layers. These concentrations also fall within the range of relevant concentrations used for transport models of biological systems such as the brain extracellular space; our layered patterning method has the potential to be extended for brain extracellular matrix modeling application in future work.^42,43^

The ability to pattern multiple materials *within the same layer* is essential to creating a structure with spatial organization in all three dimensions. Work by others to enable single layer printing of multiple gels through a multi-nozzle 3D printing system has demonstrated the interest of this capability in 3D bioprinting.^29, 44^ To demonstrate the ability to pattern materials with different compositions within a layer using our method, we built a purple agarose heart within a colorless agarose cube (Figure 3B, left). In this case, we added a simple ridge to the patterning rail to segregate the regions of different materials (purple and colorless agarose). Thus, different solutions agarose of agarose dyed different colors were allowed to flow within each layer in a defined pattern (Figure 3B, right).

A design challenge for patterning three-dimensional hydrogel structures is building unsupported features (i.e., features that cantilever into air rather than building on base material underneath). Other systems for patterning soft materials have utilized sacrificial material as a support, which is removed in a post-processing step either physically or chemically.^23, 29, 45^ The structural integrity of the desired product must be maintained after the supports are removed, which restricts the types of materials that can be used. Using our method, a dissolvable sacrificial gel approach could be employed; we also show that it is possible to create overhanging, unsupported structures without the use of sacrificial material by incrementally offsetting the patterning device from the edge of the previous layer such that each layer extends past the limits of the previous layer (Figure 4A). To exemplify this concept, the schematic in Figure 4B illustrates a rail design without an overhang (left) and with an overhang (center and right).

We found the length of overhang achievable was improved by the addition of a downward curvature to the edge of the patterning device to allow for favorable flow over the edge of the previous gel layer (Figure 4B, right). Without the added curvature, the bottom of the fluid front pins to the edge of the previous layer as expected, but the top of the advancing pre-gel solution does not continue along the patterning device (Figure 4B, center). The curvature decreases the relative ratio of the air-liquid interface to the patterning device-liquid interface; minimizing this ratio is favorable for SCF.^36^ Thus, the agarose fluid front continues along the surface of the patterning device, increasing the overhang capacity. Each patterning device has a different radius of curvature based on the curvature of the previous patterning device. Detailed descriptions of how these dimensions were determined are included in the Supporting Information and Figure S3, and the dimensions are given in Table S1. To fully demonstrate the capabilities of our method to produce geometries with overhanging features, we built an agarose cylinder with hollow winding tubes in the shape of a double helix (Figure 4C and 4D). The helix consists of 30 layers, each 500 *µ*m thick, and has a total rotation of 270 degrees.

Previously, other fabrication methods have utilized a sacrificial material to create void spaces and channels through a three-dimensional hydrogel structure^29, 38^ or have employed a casting method to create fluidic channels.^7^ Furthermore, laser photoablation can be used to create microchannels through hydrogels with high resolution,^46^ and recently multiphoton lithography has been used to create microchannels through cell-laden collagen structures.^15^ The approach we have taken here and the helix geometry proof of concept structure demonstrates the broader capacity to create gels with three-dimensional tubing and void spaces, thereby opening up the possibility of creating three-dimensional fluidic channels spanning multiple planes through a hydrogel structure without reliance on a sacrificial material or post-fabrication step.

We also demonstrate that it is possible to add open microfluidic functionality to the top of the patterning device, where the flow is also driven by capillary action.^47^ This capability can be utilized as another method by which multiple materials can be deposited in a single layer (Figure 5). We used open channels to deliver blue colored agarose to the voids in the red colored agarose layer below, which enables a single pipetting step to fill all six voids (Figure 5B).

Many materials do not have the mechanical properties necessary to create free-standing 3D structures without the use of a second support material. To overcome this challenge, we utilized the ability to pattern a single layer with multiple materials to create an agarose ‘support’ structure for unmodified type I collagen (7.5 mg/mL). Type I collagen was chosen because it is abundant in biological systems. Here, we demonstrate an alternative method for patterning layers of commercially available concentrations of unmodified collagen (Figure 6). Briefly, we patterned an agarose border on each layer, which was subsequently filled with cell-free collagen. Cell-laden collagen was then patterned over the newly established layer of collagen (Figure S4). Figure 6 shows 130 confocal image stacks stitched together of Calcein AM stained human fetal lung fibroblasts in a type I collagen structure. Cells in this structure were 87% viable, quantified using live/dead (Calcein AM/ethidium homodimer) staining (Figure S5). The ‘3’ and ‘D’ patterns represent two cell populations that were printed on two different layers, each 100 *µ*m thick and separated by a 200 *µ*m thick cell-free layer. A workflow schematic of the layer-by-layer collagen patterning process is shown in Figure S4.

For each structure presented in Figures 2-6, we used a separate patterning rail (fabricated by 3D printing or CNC milling) for each layer (Figure 7Ai). While this method is convenient for development of our method, fluid patterning by SCF can also be achieved without a physical rail by using a flat surface with patterned hydrophilic and hydrophobic regions to guide fluid flow by SCF into a desired pattern; the fluid flows along the hydrophilic tracks and pins at the hydrophilic/hydrophobic boundary (Figure 7Aii). This concept builds on past work where localized hydrophilic/hydrophobic patterning on a single flat surface was used to constrain fluids to a hydrophilic pattern.^48, 49^ Here, we extended this patterning concept to create a two-layered hydrogel structure, showing that the hydrophilic/hydrophobic patterning template can be removed after each hydrogel layer is created to iteratively build up multiple layers of hydrogel structures (Figure 7B). In the present embodiment, we plasma treated a polystyrene sheet and masked the area we wanted to preserve as hydrophilic with tape (i.e., the “W” shape shown in Figure 7Bi). A commercially available hydrophobic treatment spray was applied to create a hydrophobic background. The tape was then removed after the hydrophobic treatment, revealing a hydrophilic patterning region surrounded by a hydrophobic background. We built a two layer hydrogel (agarose) “W” structure on a polystyrene base substrate. Using stacked, glass coverslips as spacers above the polystyrene base substrate, we created a 500 *µ*m gap between the base substrate and the hydrophilic/hydrophobic patterning surface. The pre-gel solution flowed in this gap via SCF, as previously described, pinned at the interface between the hydrophilic and hydrophobic region creating the “W” pattern set by the hydrophilic region (Figure 7B-C). This result demonstrates that our open microfluidic layer-by-layer method can be implemented without a physical rail, opening up the potential for future embodiments in which the hydrophilic/hydrophobic regions are patterned via reversible or actuatable approaches.

## CONCLUSION

A novel open microfluidic method for layer-by-layer patterning of 3D hydrogel objects has been demonstrated. Our method uses fundamentally different principles—surface tension forces and open microfluidics—from existing methods to pattern hydrogel structures. For example, in contrast to patterning methods that extrude hydrogels through a nozzle, inducing strong shear force on the gel and any encapsulated cells, our method involves pipetting analogous to conventional 3D cell culture procedures. Similarly, in contrast to light-based patterning methods, our method is not limited to photopolymerizable hydrogels. Therefore, our method could potentially encapsulate cell types that otherwise would be damaged by conventional methods that rely on extrusion or chemical photoinitiators. Moreover, hydrogel waste is minimized in our layer-by-layer patterning method because it uses simple pipettes (with no dead volume) rather than requiring large vats of material or dispensing containers common in some hydrogel 3D printing methods, opening up the potential to use expensive hydrogels, rare cells such as primary patient samples, or precious reagents in small volume, a benefit of microfluidics.

We demonstrated in Figure 7 that a flat structure with a hydrophilic/hydrophobic pattern (rather than a physical rail) can be used to pattern multiple layers of hydrogel. Our hydrogel patterning method, used with surface chemistries that allow for rapid reversible patterning of hydrophilic and hydrophobic regions, could enable the creation of more automated systems to create every layer design in fractions of a second and be a step towards a novel approach to 3D bioprinting. Additionally, the processing time could be improved in a future embodiment that provides tighter local temperature control, affecting faster gelling. Importantly, using the embodiment shown in the present manuscript, we demonstrated layered patterning of cell-laden structures using standard cell culture materials and procedures without subjecting the fluid to external pressure, suggesting that this method has the potential to be extended for use with other cell types. Our capacity to control the location of different gel materials within and between layers suggests the possibility to create 3D organoid models by directing the patterning of cell types and culture materials in 3D space. Building open microfluidic channels directly into the patterning device (Figure 5) can enable efficient delivery of reagents, cells, or other components into void spaces with a single dispensing step. Finally, as demonstrated, our method can be used to create overhanging structures and hollow tubes, extending the capability of adding fluidics (channels, voids, etc.) throughout the gel itself, thus opening up possibilities for complex three-dimensional models and systems.

## Supporting information

Figure 6 original image file

Void filling Figure 5 video

Hexagon and Flower Figure 2B video

## ACKNOWLEDGEMENTS

We thank Ivor Clinton for assistance with photography, Jing Lee for video editing, and Dr. Mary Regier for helpful discussions. We gratefully acknowledge funding from NIH 1R01HD090660-01A1, the Arnold and Mabel Beckman Foundation (Beckman Young Investigator Award), and the University of Washington. Research reported in this publication was supported in part by the Eunice Kennedy Shriver National Institute Of Child Health & Human Development of the National Institutes of Health. The content is solely the responsibility of the authors and does not necessarily represent the official views of the National Institutes of Health.

