## Supplementary figures and images for "Layer-by-Layer Fabrication of 3D Hydrogel Structures Using Open Microfluidics"

### Figure 6 original image file

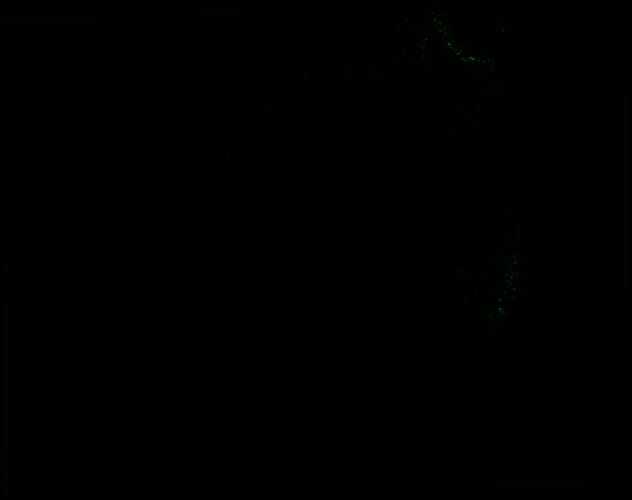
