## Supporting Information for "Layer-by-Layer Fabrication of 3D Hydrogel Structures Using Open Microfluidics"

### Supporting Information Contents

Extended Methods and Materials
Figure S1. Details of patterning device design.
Figure S2. Engineering schematics of patterning devices in Figure 2B.
Figure S3. Schematic of helix and design parameters for unsupported overhangs.
Figure S4. Schematic of workflow for building cell-laden type I collagen structure.
Figure S5. Live/dead image of cells in Figure 6.
Figure S6. Photo of patterning setup in Figure 7.
Table S1. Calculated designed curvature values for patterning devices in Figure 4.
Video of formation of hexagon and flower from Figure 2B.
Video of multiple voids filling from Figure 5.
Video of 'W' patterning from Figure 7.
Original (.tif) files for confocal imaging in Figure 6.
CAD files (.sldprt) of patterning devices used in Figure 2B and template used to design all patterning devices – please contact corresponding author for files.

### Materials and Methods

#### Derivation for SCF condition in rail-based microchannels

In the case where a channel is comprised of multiple surfaces with different materials and unique contact angles, the condition for advancement of a fluid in the channel is given by a condition on the Cassie Angle, $\theta^*$(1a), referred to as the Generalized Cassie law (1b), where $f_i$ is the fraction of the channel's cross-sectional perimeter composed of a material $i$, and $\theta_i$ is the contact angle of material $i$.

$$\cos(\theta^*) = \sum_i f_i \cos(\theta_i) \quad \text{(Eq. 1a)}$$

$$\cos(\theta^*) > 0 \quad \text{(Eq. 1b)}$$

Given the cross-sectional geometry of a rail-based channel and the convention that the contact angle of fluid on air is 90°, Eq. 1 can be rewritten as:

(Eq. 2)

Where $w_1$ is the width of the channel ceiling, $w_2$ is the width of the channel floor, and $h$ is the distance between the channel ceiling and the floor. Assuming that $w_1 = w_2$, Eq. 2 reduces down to:

$$\frac{h}{w_1} < \frac{\cos(\theta_1) + \cos(\theta_2)}{2} \qquad \text{(Eq. 3)}$$

where $h$ is the height of the patterned layer (defined by the distance between the patterning area of the patterning device and the underlying substrate), $\theta_1$ is the contact angle of the patterning device, and $\theta_2$ is the contact angle of the underlying substrate.

**Design of hollow double helix**

The helix consists of a total of 30 layers of agarose (including the base layer), which were all patterned via SCF and rail-based technology. Each layer looks like a button with two holes, that is rotated about the center for each subsequent layer. Figure S3A illustrates a top-down schematic for the helix design. The length, $d$, is the diameter of the holes, which is 3 mm for all layers on both sides. $r$ is the distance from the center of the design to the center of the holes on both sides, and for all layers is 2.5 mm. The angle of rotation about the center of the design is denoted by $\phi$, and the maximum overhang length is denoted by $s$. For the first two layers, the angle of rotation, $\phi$, is 5° with a designed overhang, $s$, of 240 $\mu$m to establish the overhang. After the overhang was established, the remaining layers were rotated 10° for a total designed maximum overhang of 479 $\mu$m. A new device was used for each layer with an alignment marker for consistency in the degree of rotation. The helix was designed to include a curvature that outlined the two holes which allowed for SCF to continue to the edge of the desired overhang. These values can be found in Table S1.

**Imaging of hollow double helix**

A Nikon DSLR camera was used to take images of the double helix. The helices were removed from the well plate with a spatula and placed on a standard No. 1.5, 25 mm square coverslip (Fisherbrand). A 1 mL syringe with a 18 gauge needle with 18 gauge PTFE tubing attached to reach the bottom of the helices was used to load the helices with dye. A syringe with a 25 gauge needle was used to remove any air pockets that formed from filling the hollow tubes with dye. During loading, the tubing damaged the walls of the agarose and these irregularities filled with dye (Figure 4C, right). Yellow and blue India ink dye (Dr. Ph. Martin's Bombay India Ink) was used to fill the helix (Figure 4C, right), and dextran with fluorescein 70,000 MW in DI water at 10 mg/mL (Invitrogen) was used to fill a separate helix in Figure 2C. To take the photo in Figure 2C, the fluorescent dye was excited using a 365 nm lamp (Spectroline). Helix images were processed using Adobe Photoshop CC 2018 using a uniform brightness/contrasts adjustment.

**Imaging and design of overhanging cross-sections**

The cross-section images shown in Figure 4A were designed such that the overhang and curvature of each layer corresponded to the maximum overhang in the first seven layers of the helix. The curvature information can be found in Table S1 where the first seven layers correspond to the overhang cross-section device, as well as the first seven layers of the helix. To image the cross-section, a smooth edge was cut using a coverslip (Fisherbrand) and the overhang was transferred to another coverslip for imaging with a DSA25E goniometer (Kruss). The images in Figure 4A are separately constructed agarose structures, cut and imaged at different stages.

**Design of curvature for overhanging features**

In order to achieve spontaneous capillary flow for all layers of the seven layered overhanging structure (Figure 4A) and the hollow double helix structure (Figure 4C, Figure 2C), the geometry of curvature was adjusted for each layer. If each layer had the same curvature, then the maximum distance (air gap) between the two layers would become great enough that spontaneous capillary flow is inhibited. This concept is illustrated in Figure S3C, as denoted by the red arrow. However, by applying a calculation to change the curvature dimensions of the current layer based upon the dimensions of the previous layer, the distance between layers can be minimized allowing for SCF, illustrated by the reduction of the double-headed red arrow length (Figure S3C). Note that the overhang dimensions for the seven layered overhanging structure in Figure 4A are the same as the dimensions for the first seven layers of the helix. The calculation was made as follows: for the first overhanging layer, layer 2, the radius of curvature is set to 1 mm. Every subsequent layer's radius of curvature is increased by 0.5 mm, i.e., layer 3 is 1.5 mm, layer 4 is 2 mm, etc. Take any layer, $n$ where $n > 1$, and the radius of curvature, $r_n$ is then given by Eq. 4.

$$r_n = 0.5n \quad \text{(Eq. 4)}$$

Layer 1 has no radius of curvature, as it is not an overhanging layer. The curvature of layer $n$ was designed such that the curvature began at half the distance of the x-component of the previous layer, $x_{n-1}$, which is illustrated in the diagram below.

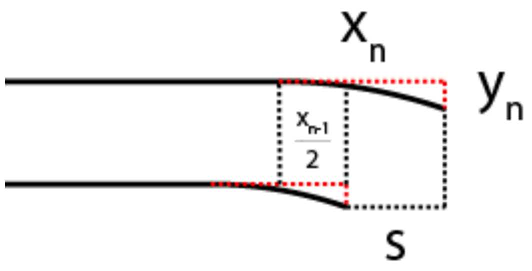


Note that $s$ is the designed maximum overhang, which for the helix design is also denoted by $s$ in Figure S3A. The $s$ component of the curvature for layer $n$, $x_n$, is then given by taking one half of the x-component of the previous layer, and adding this to the maximum overhang, $s$, described in Eq. 5.

$$x_n = \frac{x_{n-1}}{2} + s \quad \text{(Eq. 5)}$$

Note that the maximum overhang, $s$ is set by the angle of rotation for the helix ($\phi$ in Figure S3A). Using the calculated x-component and the radius of curvature for layer $n$, the y-component of the curvature can then be derived. The schematic below shows the geometrical schematic to calculate these layers.

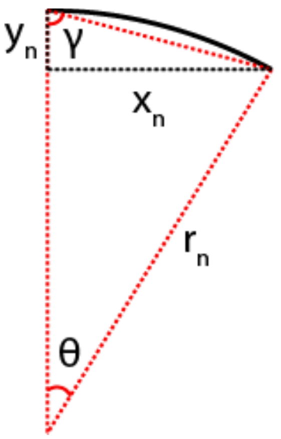


To calculate the angle, $\theta$, which represents the arc angle of the circle that encompasses the curvature, the geometric relation of the radius of curvature, $r_n$, to the x-component of the curvature was used, and described in Eq. 6 where $\theta$ is in degrees.

$$\theta = \arcsin\left(\frac{x_n}{r_n}\right) \quad \text{(Eq. 6)}$$

Using this angle and the fact that the red dotted triangle is an isosceles triangle, the value of angle $\gamma$ could be calculated using Eq. 7, where $\gamma$ and $\theta$ are in degrees.

$$\gamma = \frac{180 - \theta}{2} \quad \text{(Eq. 7)}$$

Then, the value of $y_n$ can be calculated using Eq. 8.

$$y_n = \frac{x_n}{\tan(\gamma)} \quad \text{(Eq. 8)}$$

These calculations were applied to all layers to derive the values in Table S1. Note that the first overhanging layer, layer 2, was set such that the curvature began at the edge of layer 1, necessitating that the x-component was set to 239.9 $\mu$m.

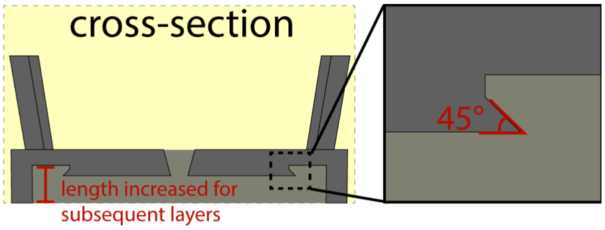


**Figure S1.** Cross-section of patterning device with inset of 45° angled edge to prevent capillary rise. If the patterning device contained a simple 90° angled edge rather than the 45° angled edge shown here, the pre-gel solution would wet the vertical edge of the patterning device due to capillary rise. The diagram also shows the portion of patterning device that is adjusted to increase the height of the patterning area for subsequent layers of hydrogel.

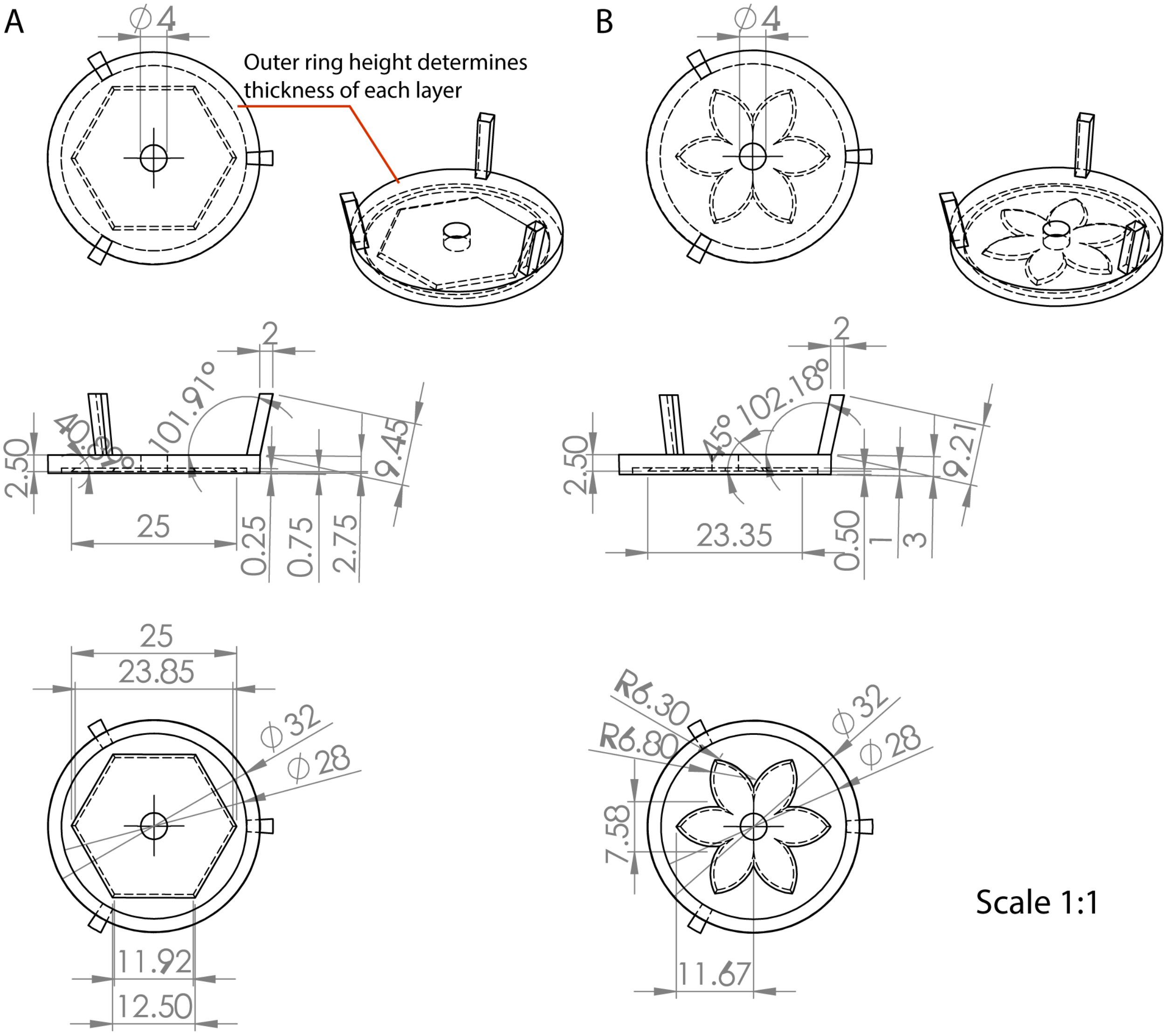


**Figure S2.** Schematic diagrams illustrating the dimensions of the patterning devices in Figure 2B. All dimensions are in mm. CAD files are also included. (A) First layer hexagon pattern. Red lines indicate the ring ('foot' of the device) that sits flush to the well plate floor which is adjusted for each patterning device to determine layer thickness. (B) Second layer flower pattern.

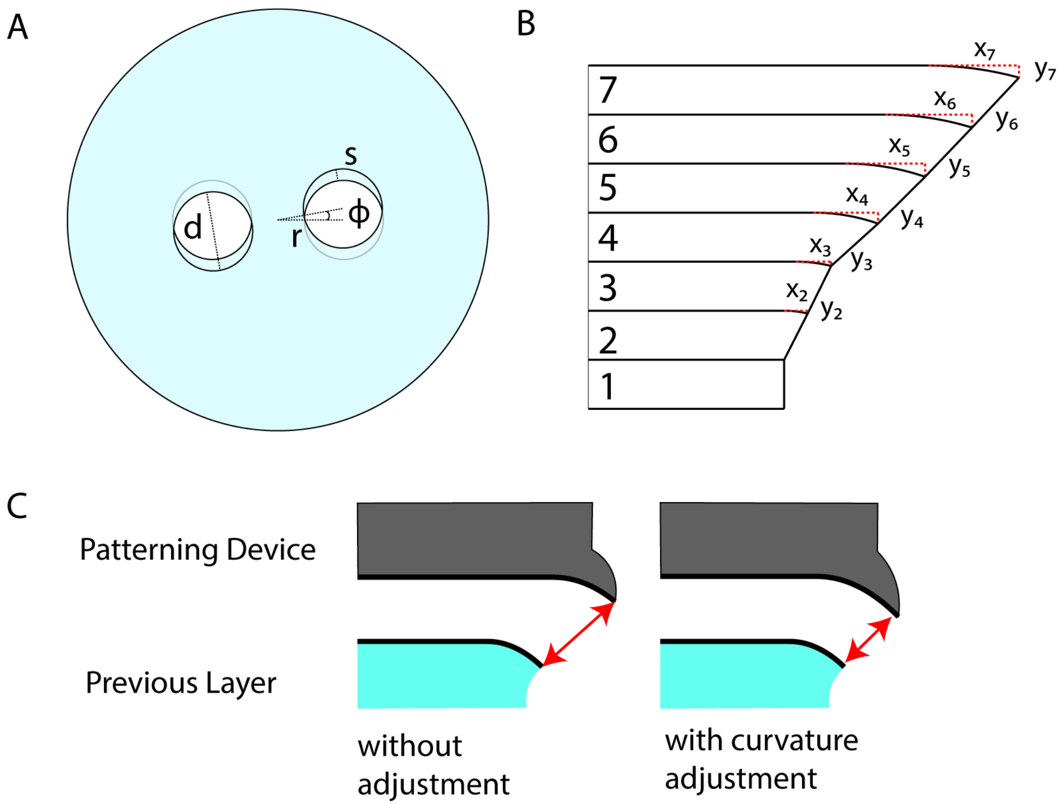


**Figure S3.** Schematic of helix and overhang design used to make the structures shown in Figure 4 of the manuscript. (A) Dimensions of each design parameter where $d$ is diameter of hole, $r$ is distance from center of the design to the center of the hole, $\phi$ is the angle of rotation from layer to layer, and $s$ is the maximum overhang length. (B) Schematic of the overhang design for 7 layers, showing the curvatures for each layer. Note: there is no curvature for the first layer. (C) The effect of adjusting the curvature based on the previous curvature on the distance between the two layers, as denoted by the red arrows, which limits the surface capillary flow to the edge of the overhang. The left diagram shows the same curvature for both layers and the right diagram shows curvature adjustment on the patterning device based on the previous layer.

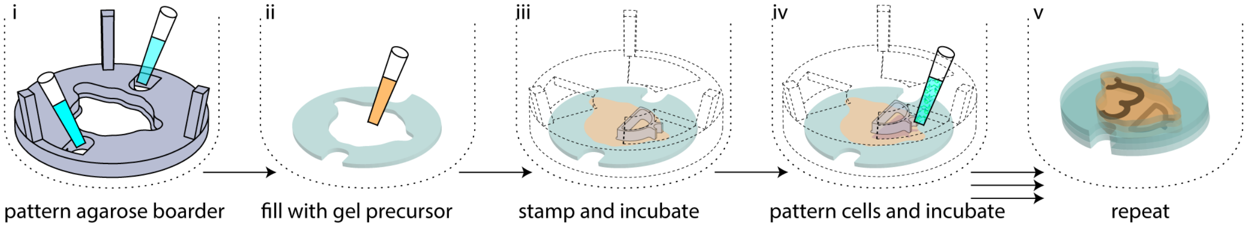


**Figure S4.** Schematic workflow of agarose and collagen patterning process used to make the cell-laden collagen structure structure in Figure 6. (i) A 200 $\mu$m tall agarose border is patterned in a 6-well plate. (ii) The agarose border is then filled with collagen gel precursor. (iii) A 200 $\mu$m tall 'D' patterning device is then placed in the well, which precisely sets the height of the collagen fill to 200 $\mu$m (the patterning device is shown as two parts for visualization purposes, where the main device component is shown in dotted lines and the 'D' patterning component is shown in transparent grey). The well plate is then incubated with the device in place for 10 minutes to polymerize the collagen. (iv) The previous patterning device is then removed, and an identical, 300 $\mu$m tall patterning device is placed in the same position. 8 $\mu$L of cell-laden collagen gel precursor solution is then pipetted into the loading region of the patterning device, at which point the gel precursor solution flows through the 'D' pattern. These 4 steps are repeated once more with a '3' pattern, and then again with no pattern to yield (v) the final structure. As described in the manuscript, the final structure consists of a first cell-free layer that is 200 $\mu$m tall, then a cell-laden collagen layer containing the 'D' pattern that is 100 $\mu$m tall, then a cell-free layer that is 200 $\mu$m tall, then a finally cell-laden collagen '3' pattern that is 100 $\mu$m tall, then a cell-free layer.

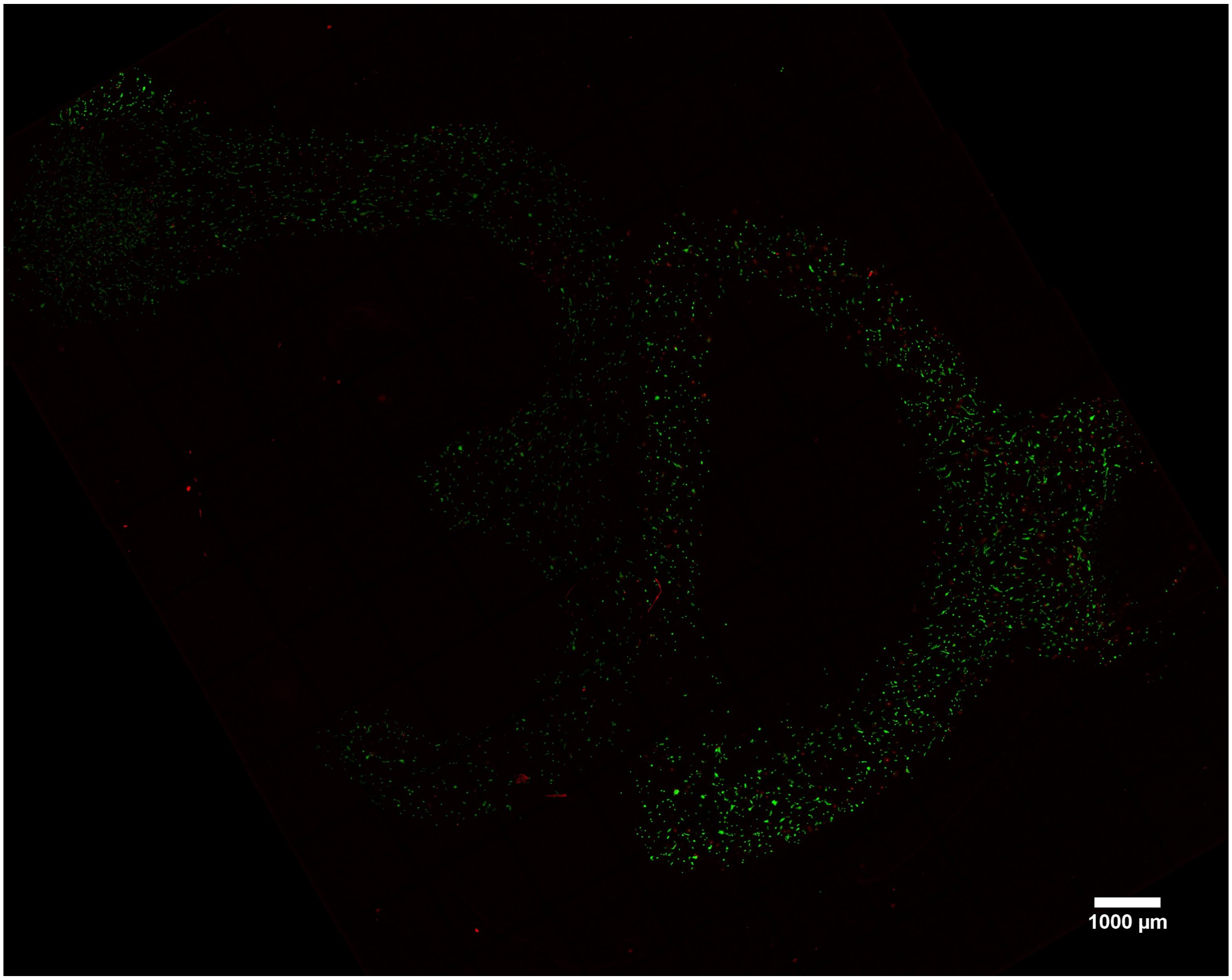


**Figure S5.** Confocal images showing live/dead staining of human fetal lung fibroblast cells in 3D collagen structure from Figure 6. Image comprises 130 fields of view (which each contain 27 Z-stacked images) stitched together. Green is Calcein AM, which stains the cytoplasm of live cells, and red is ethidium homodimer, which stains the nuclei of dead cells. Viability (87%) was quantified in FIJI image processing software (using a minimum cell area of 25 $px^2$ for the live and dead stains and a minimum circularity of 0.25 for the dead stain).

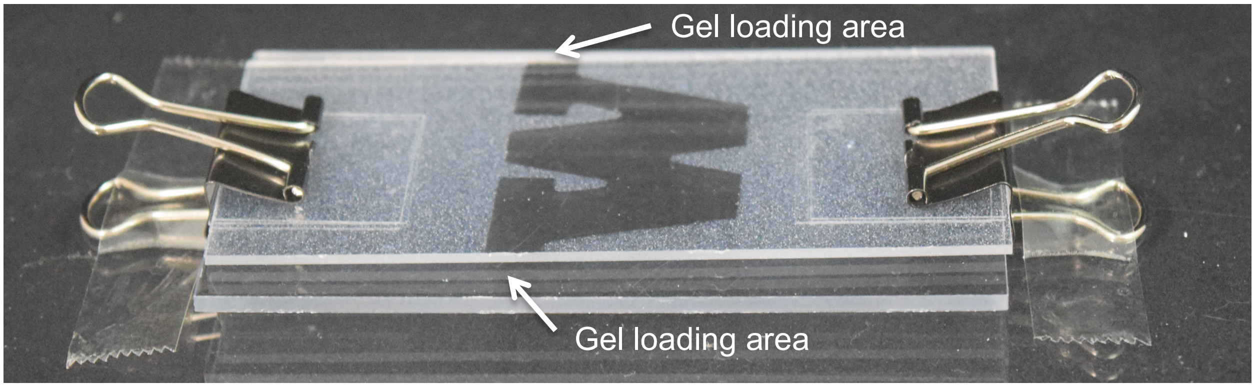


**Figure S6.** Set up of Figure 7Aii before pre-gel solution is pipetted into gel loading areas.

**Table S1.** Calculated designed curvature values for all patterning devices used for double helix structure.*

| Layer | Rotation angle, ϕ (degree)** | Maximum overhang*** ($\mu$m) | Radius of curvature (mm) | Horizontal length of curvature, x ($\mu$m) | Vertical length of curvature, y ($\mu$m) |
|---|---|---|---|---|---|
| 1 | 0 | 0 | 0 | 0 | 0 |
| 2 | 5 | 239.9 | 1.0 | 239.9 | 29.2 |
| 3 | 5 | 239.9 | 1.5 | 359.9 | 43.8 |
| 4 | 10 | 479.4 | 2.0 | 659.3 | 111.8 |
| 5 | 10 | 479.4 | 2.5 | 809.1 | 134.5 |
| 6 | 10 | 479.4 | 3.0 | 884.0 | 133.2 |
| 7 | 10 | 479.4 | 3.5 | 921.4 | 123.5 |
| 8 | 10 | 479.4 | 4.0 | 940.1 | 112.0 |
| 9 | 10 | 479.4 | 4.5 | 949.5 | 101.3 |
| 10 | 10 | 479.4 | 5.0 | 954.2 | 91.9 |
| 11 | 10 | 479.4 | 5.5 | 956.5 | 83.8 |
| 12 | 10 | 479.4 | 6.0 | 957.7 | 76.9 |
| 13 | 10 | 479.4 | 6.5 | 958.3 | 71.0 |
| 14 | 10 | 479.4 | 7.0 | 958.6 | 65.9 |
| 15 | 10 | 479.4 | 7.5 | 958.7 | 61.5 |
| 16 | 10 | 479.4 | 8.0 | 958.7 | 57.7 |
| 17 | 10 | 479.4 | 8.5 | 958.7 | 54.2 |
| 18 | 10 | 479.4 | 9.0 | 958.7 | 51.2 |
| 19 | 10 | 479.4 | 9.5 | 958.7 | 48.5 |
| 20 | 10 | 479.4 | 10.0 | 958.7 | 46.1 |
| 21 | 10 | 479.4 | 10.5 | 958.7 | 43.9 |
| 22 | 10 | 479.4 | 11.0 | 958.7 | 41.9 |
| 23 | 10 | 479.4 | 11.5 | 958.7 | 40.0 |
| 24 | 10 | 479.4 | 12.0 | 958.7 | 38.4 |
| 25 | 10 | 479.4 | 12.5 | 958.7 | 36.8 |
| 26 | 10 | 479.4 | 13.0 | 958.7 | 35.4 |
| 27 | 10 | 479.4 | 13.5 | 958.7 | 34.1 |
| 28 | 10 | 479.4 | 14.0 | 958.7 | 32.9 |
| 29 | 10 | 479.4 | 14.5 | 958.7 | 31.7 |

*First seven layers curvature and overhang information also corresponds to the cross-section device in Figure 4A.
**Rotation angle represents the rotation change from the previous layer. Values are in degrees.
***Maximum overhang corresponds to $s$ in Figure S3A.
